# Complexity and 1/f slope jointly reflect brain states

**DOI:** 10.1101/2020.09.15.298497

**Authors:** Vicente Medel, Martín Irani, Nicolás Crossley, Tomás Ossandón, Gonzalo Boncompte

## Abstract

Characterization of brain states is essential for understanding its functioning in the absence of external stimuli. Brain states differ on their balance between excitation and inhibition, and on the diversity of their activity patterns. These can be respectively indexed by 1/f slope and Lempel-Ziv Complexity (LZc). However, whether and how these two brain state properties relate remain elusive. Here we analyzed the relation between 1/f slope and LZc with two in-silico approaches and in both rat EEG and monkey ECoG data. We contrasted resting state with propofol anesthesia, which directly modulates the excitation-inhibition balance. We found convergent results among simulated and empirical data, showing a strong, inverse and non trivial monotonic relation between 1/f slope and complexity, consistent at both ECoG and EEG scales. We hypothesize that differentially entropic regimes could underlie the link between the excitation-inhibition balance and the vastness of the repertoire of brain systems.

## Introduction

Spontaneously occurring brain activity patterns in the cerebral cortex constitute the so-called brain states (1, 2). These are present without a direct link to external stimuli, and constitute the basis of essential cognitive processes like attention (3–5) and global states of consciousness (GSC), such as sleep, wakefulness and anesthesia (6, 7). One of the most prominent strategies to characterize brain states has been to analyze the spectral properties of their associated field potentials like electroencephalogram (EEG) and local field potential (LFP). In the particular case of attention, it has been shown that both induced and spontaneous modulations of properties of alpha-band oscillations broadly explain the attentional state of subjects (8–10). However, the characterization of GSC in terms of the unique neural properties of their associated brain states has proven to be more elusive.

Spectral characteristics of brain field potentials cannot fully distinguish between GSC (11). This is well illustrated by the case of anesthetics that equally produce a cease of phenomenological experiences in loss of consciousness, but show diverse spectral signatures. For example, transitions from wakefulness to anesthesia induced by propofol increase and frontalize alpha oscillations (8 - 12 Hz), whereas dexmedetomidine anesthesia instead induces spindle-like activity (12 - 15 Hz) (12, 13). A broader approach considers oscillatory activity as only one aspect of neural activity, given that another important source of variance in neural recordings can be captured by aperiodic neural activity with complex neural dynamics (14, 15). From this perspective, our understanding of GSC and its relation to brain states remains limited partly due to the canonical focus on narrow-band oscillations, which marginalizes non-linear activity to the status of ‘background’ activity or irrelevant ‘noise’. However, this background noise may contain crucial information to bridge the gap between brain states and GSC.

Cortical neurons in awake animals show strong membrane potential fluctuations generating irregular discharges, known as high conductance states (16). These states generate the background activity that supports high-order processes. It has been shown that neurons can achieve irregular firing patterns with balanced excitatory and inhibitory synaptic activity (17, 18). From this perspective, brain states depend on global brain variables, such as relative levels of excitation and inhibition (19). Moreover, computational characterizations of the balance between excitation and inhibition (E/I balance), from local circuit activity to whole-brain modeling, have shown its relevance on modulating information transmission and entropy (20–22). On the other hand, perturbations to the E/I balance have shown to be related to pathological brain activity (23) and neuropsychiatric disorders (19, 24–26). A proposed way to quantify E/I balance is the slope of the power law decay of spectral power of brain field potentials. Specifically, models have shown that the background 1/f slope of the power spectral density (PSD) can emerge from the sum of stochastic excitatory and inhibitory currents (27–29). Moreover, empirical validation of these models has shown that the E/I balance can be inferred from background activity by parameterizing the 1/f shape of the PSD (29, 30).

Interest in the detailed informational structure of brain states has produced a recent surge of information theory-based approaches (31–33). Data analysis strategies based on Lempel-Ziv complexity (LZc; (34)), like the Perturbational Complexity Index (35) have been successful for characterizing subject’s GSC during dreamless sleep and during anesthesia-induced unconsciousness, with partial independence of the anesthetic used. It has been shown that LZc normally decreases concomitantly with the loss of phenomenological possibilities (e.g. (36, 37)), which is consistent with theoretical views of consciousness (38, 39). Lempel-Ziv complexity algorithm computes the number of non-redundant patterns of a signal (34), which in turn, when applied to brain data, is related to the diversity of the repertoire of brain activity patterns (40). During the transition from wakefulness to sleep or anesthesia, the number of possible experiences and cognitive processes that one can have is greatly reduced. Thus, it is expected that the complexity of brain activity follows the same pattern. In fact, this reduction of the repertoire of brain activity has been seen in rats at the single neuron level using a myriad of convergent measures of cortical diversity, including LZc (41), suggesting that LZc can be applied as a multiscale proxy of neural repertoire.

Although 1/f slope and LZc reflect different brain state properties and have distant theoretical origins, one coming from spectral analysis and the other from Information Theory, both have been separately shown to correlate with GSC (33, 42). We hypothesize that this could be due to an underlying intrinsic relation between E/I balance and the repertoire of activity patterns in brain systems. Here we employed four complementary approaches to study the relation between 1/f slope and LZc and thus implicitly between E/I balance and the abundance of non-redundant repertoire in brain field potentials. We analyzed this relation in: (1) a simple inverse Discrete Fourier Transform (iDFT) model, (2) a biophysical model, (3) rat EEG, and (4) monkey ECoG during wakefulness and propofol anesthesia. Our results consistently show an inverse and non-trivial relation between 1/f slope and LZc in brain field potentials, suggesting that both could be related to the underlying entropic state of cortical systems.

## Results

### iDFT model show a robust and specific relation between LZc and 1/f slope

In order to analyze the relation between the spectral power-law slope and LZc we began by constructing, from the frequency domain, sets of time series with different 1/f spectral power decay slopes (Figure 1A; see Materials and Methods). We simulated 256 time series with slopes ranging from 0 to 2, and calculated LZc for each one. Figure 1B illustrates that, for these simple iDFT models, 1/f and LZc follow a strict monotonically descending behavior, with lesser complexity values for time series with a steeper slope. This general behavior is to be expected: slopes near zero reflect white noise (high LZc), while on the other hand high slopes reflect time series with significant power only in low frequencies (periodic signals with low LZc). Interestingly, we found that LZc had a one to one mapping with 1/f slope. This relation can be robustly adjusted (*R*^2^ > 0.99) to an x-inverted asymmetrical sigmoid function (see Materials and Methods).

**Fig. 1.**
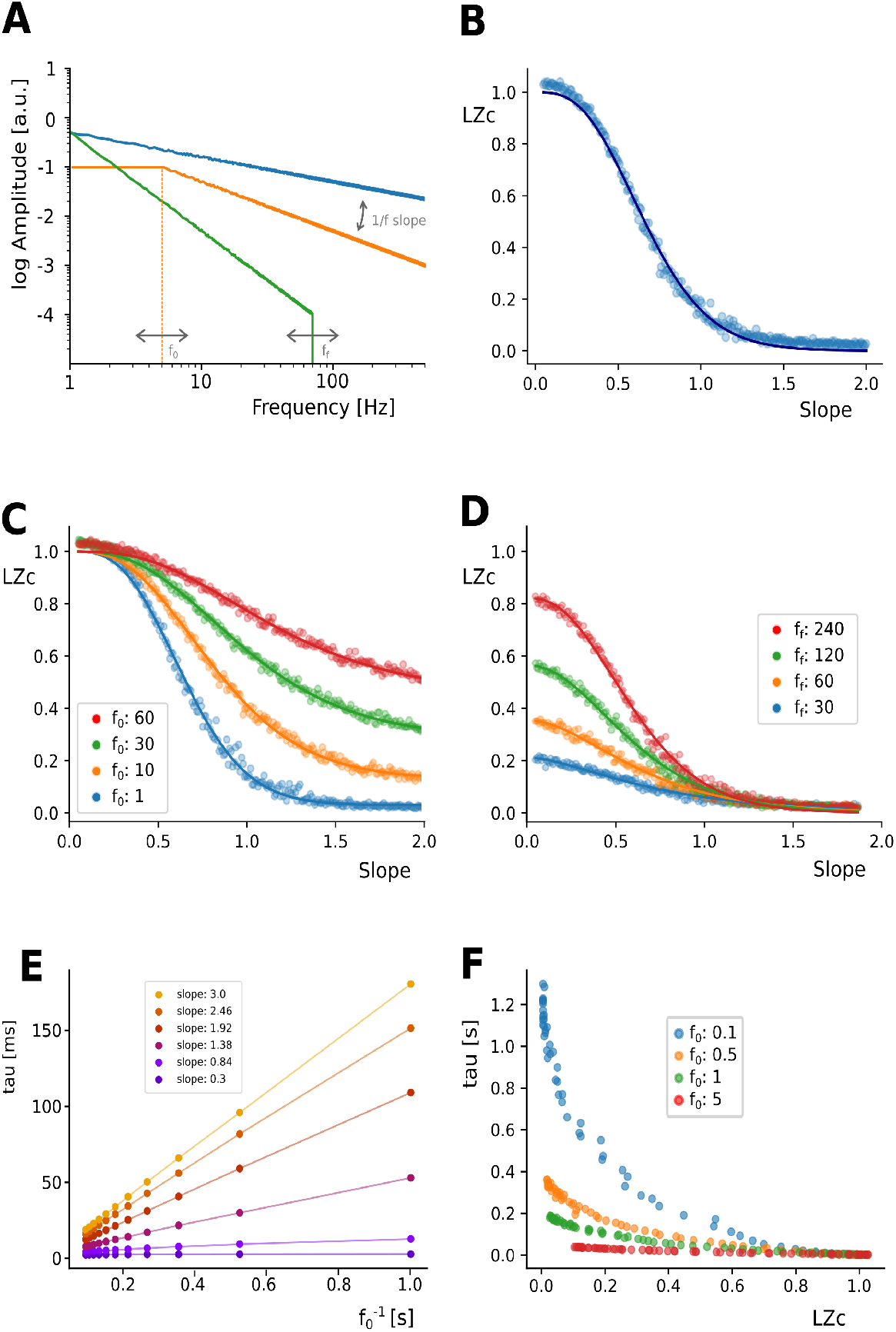
iDFT model illustrates a strict relation between LZc and 1/f slope. (**A**) Depiction of the parameters employed. Time series were constructed by first defining a power-spectrum structure according to power law decays with various slopes, initial frequencies (*f_0_*) and final frequencies (*f_f_*). Afterwards this was converted to the time domain by means of iDFT. (**B**) Scatterplot showing the relation between LZc and 1/f slope. (**C**) Scatterplot depicting how the relation between LZc and slope changes with different initial frequencies of the power law. (**D**) Same as C but using various final frequencies. Solid lines in B, C and D correspond to best fit of eq 3-4 (**E**) Scatterplots depicting the strong and linear relation that exists between timescale (tau) and the period of the slower oscillation of the power-law behaviour, in other words (the inverse of *f*_0_) for different slope values. (**F**) Scatteplot showing the relation that exists between LZc and tau for different values of initial frequency *f_0_*.

Electrophysiological field potential signals (e.g. EEG and ECoG) have been shown to present only partial power-law behavior (43). In other words, only part of their spectrum follows a clear spectral power-law distribution. In an attempt to emulate this, we introduced two types of constraints to the spectral construction of signals: an initial (*f_0_* also referred to as “knee”) and a final (*f_f_*) 1/f frequency (see Materials and Methods). Both constraints are illustrated in Figure 1A. We found that the introduction of greater f0 values (Figure 1C) generated signals with greater complexity across all slopes tested. This effect was more prominent for higher slope values than for lower slopes. Interestingly, the introduction of *f_0_* higher than 1Hz reduced the dynamical range of the observed LZc (no longer ranging from 0 to 1). On the other hand a final frequency ff (homologous to a low-pass filter), reduced the complexity of the resultant time series (Figure 1D). This effect was more marked in signals with lower slope values. Similarly to *f*_0_, we found that ff also reduced the dynamical range of possible complexity values, but in a different way: LZc now ranged from zero to a value lower than 1. Regardless of these spectral constraints, we found that the slope vs. LZc relationship could be modeled with a simple set of related equations (see Materials and Methods, equations 3 and 4), with a robust goodness of fit (all *R*^2^ > 0.98, see Supplementary Materials).

Given this robust relation between the general spectral properties of synthetic signals and their LZc, we asked whether this relation could also be tracked by another non-linear characteristic of time series, namely their autocorrelation function (ACF). This was also motivated by evidence suggesting that ACF’s tau (also called timescale) can be obtained based on the initial (or knee) frequency of the power-law decay in neural data (44). To evaluate this, we constructed new sets of iDFT time series and calculated tau for each one of them. As expected, we found that increasing slopes generally produced longer timescales. However, this relation was strongly dependent on *f*_0_ (and its inverse;Figure 1E). We consistently found that the period of the initial frequency 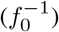 had a linear relation with tau, and that the slope of this relation (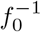 vs tau) was dependent on the spectral 1/f slope. Thus, the relationship between tau and 1/f slope is strongly modulated by the initial frequency (f0) of the corresponding spectral power-law behaviour. In the same line, we next explored the possible relation between tau and LZc by constructing iDFT signals that varied in 1/f slope and *f*_0_. As expected, we found that generally tau presented an inverse relation with LZc, showing that faster timescales were associated with higher complexity. However, this relation was almost completely dominated by 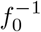. Figure 1F shows that, while for small 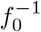 values (e.g. 0.1 Hz) tau and LZc are notoriously related, this relation is severely distorted and diminished when higher, more physiological values of 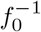 are employed (e.g. > 2Hz). This shows that, although in some circumstances tau is robustly related to 1/f slope and LZc, this relation is dominated by the initial frequency of the spectral power-law decay. Thus, tau does not seem to fully capture the richness of the spectral structure of time series.

### Conductance-based neural network model and kernel-based LFP estimation

The spectral 1/f slope has been suggested as a proxy of the background state and the balance between excitation and inhibition in cortical circuits (15, 27, 29, 30, 45). In this line, we hypothesized that E/I balance could also be related to the repertoire of brain activity as indexed by LZc. To examine this, and to test whether iDFT model predictions are also present in biologically-based model settings, we use a recurrently-connected network of spiking neurons with conductance-based synapses. The network is composed of 80% AMPA-like excitatory and 20% GABA-like inhibitory neurons. All neurons were modeled as leaky integrate-and-fire (LIF) receiving two types of inputs: a sensory-driven thalamic input and an intracortical input (30, 46). From this model, we computed the network’s local field potential (LFP) by fitting a kernel function defined by the unitary contribution of the spiking of neurons to the LFP (47). In this simulation, we manipulated E/I ratio by emulating different GABAergic tones; we introduced a scaling factor that multiplied unitary GABA-like conductances (see Materials and Methods. This manipulation robustly and systematically modulated the PSD structure (Figure 2C) and the firing rate of both excitatory and inhibitory populations (Figure 2D). Consistent with previous reports (29, 30), we found that PSD displays a modulation of its aperiodic features at the approximate frequency range of 30-70 Hz (Figure 2C). We found that increasing the scaling factor (that is, E/I ratio is shifted towards inhibition) was consistently related with both an increase in the 1/f slope (Figure 2E) and with a decrease in LZc (Figure 2F). (Figure 2G) summarises both these effects by showcasing the strong and inverse relation that exists between LZc and 1/f slope across E/I balances using this modelling strategy (LZc vs 1/f slope: Spearman rho: 0.96, p<0.001). These results indicate that, in a biophysically inspired, recurrently-connected neural network of excitatory and inhibitory neuronal populations, changes in the synaptic E/I conductances could be simultaneously inferred from LFP features such as LZc and 1/f slope.

**Fig. 2.**
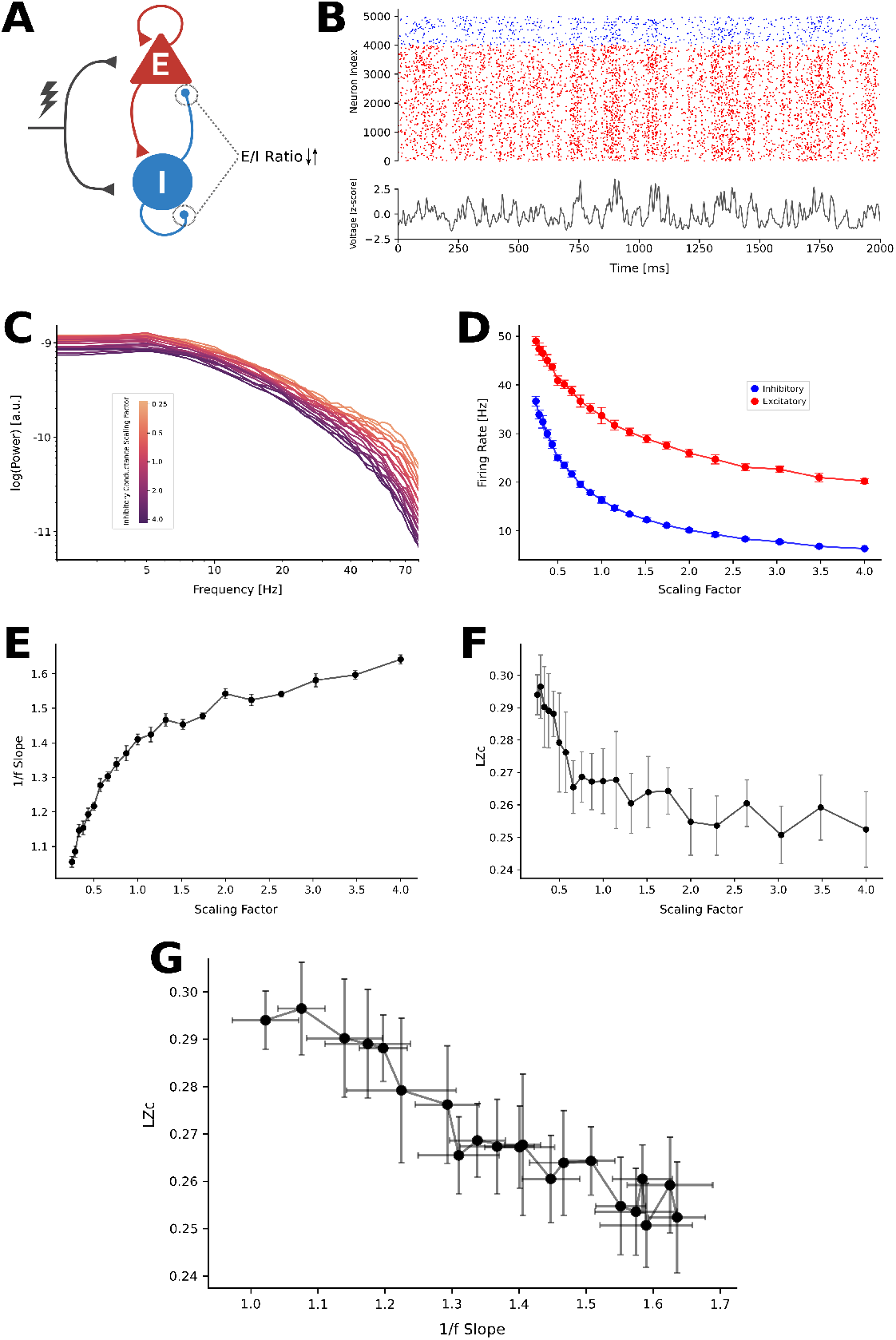
1/f slope and LZc present an inverse relation across E/I balances in a recurrently connected LIF model. (**A**) Schamatic of the model structure. Excitatory (red) and inhibitory (blue) neurons are externally stimulated by thalamic inputs (lightning) and by cortico-cortical connections. E/I balance was manipulated by changing both inhibitory-inhibitory and inhibitory-excitatory conductandes. (**B**) Top) Representative rasterplot depicting spikes across time for excitatory (red) and inhibitory(blue) simulated neurons. Bottom) Representative LFP calculated from spiking activity. (**C**) PSD of simualted signals with different scaling factors. Bigger scaling factors imply more inhibitory activity. (**D**) Average firing rate across scaling factors for both inhibitory (blue) and excitatory (red) neurons. (**E**) 1/f slope as a function of scaling factor. (**F**) LZc as a function of scaling factor. (**G**) Scatterplot showing the strong and inverse relation between LZc and 1/f slope for different E/I balances (LZc vs 1/f slope: Spearman rho: 0.96, p<0.001).

### Rat EEG and Monkey ECoG

Next, we asked whether the impact of modifying E/I balance on the relationship between 1/f slope and LZc seen in our model could be reproduced in empirical electrophysiological data. We analyzed two open datasets, both obtained during resting-state and during increased cortical inhibition by propofol: a macaque monkey ECoG (48), and an epidural EEG in rats (49, 50).

Propofol is known to directly enhance GABAergic inhibitory activity, and thus reduce E/I balance (51). In accordance with our previous results, we observed an increase of the 1/f slope for propofol (conscious state main effect’s F(1) = 1034, p < 0.001, η^2^ = 0.467; simple main effects (awake vs. anesthesia) for all monkeys (p < 0.001)) and reduced LZc with respect to wakefulness (conscious state main effect F(1) = 442, p < 0.001, η^2^ = 0.063; simple main effects (awake vs. anesthesia), p < 0.001) and rat EEG (p < 0.0001). This is illustrated in Figure 3 C-F.

**Fig. 3.**
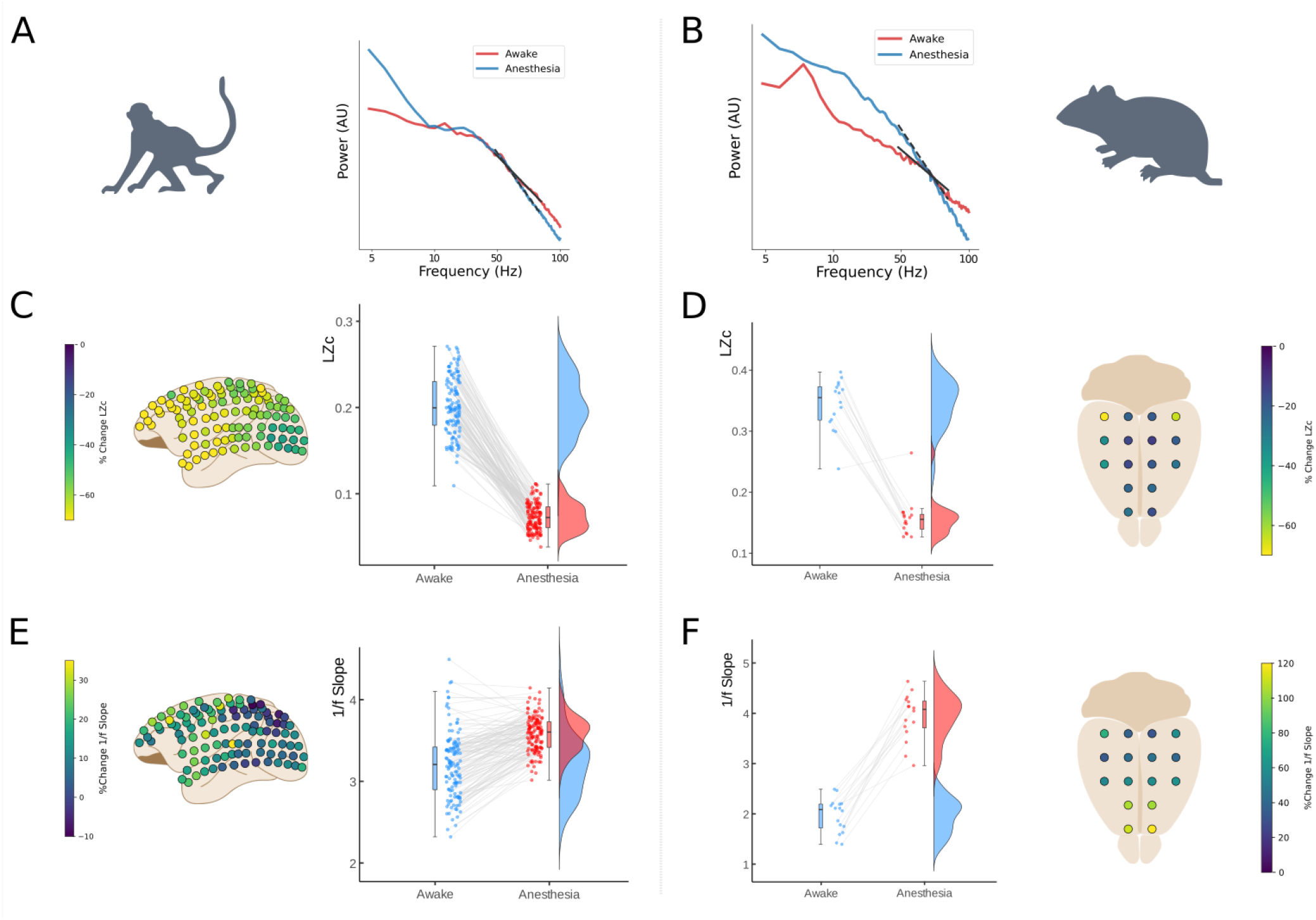
Title of Figure. (**A-B**) Power spectral Density with 1/f slope fittings (30-80 Hz range) of representative macacque ECoG (Left, Monkey 1) and rat EEG (Right, Rat 1) in Awake (red) and propofol anesthesia (blue) brain states. (**C-D**) In borders, the topography of the percentage of change from baseline for Lempel-Ziv complexity for macacque (border left) and rat (border right). In middle, a paired-sample raincloud plot showing LZc individual electrode changes across brain states for macacque (middle left) and rat (middle right). (**E-F**) In borders, the topography of the percentage of change from baseline for 1/f slope for macacque (border left) and rat (border right). In middle, a paired-sample raincloud plot showing 1/f slope individual electrode modulation across brain states for macacque (middle left) and rat (middle right).

To estimate the spatial correspondence of the effect of propofol, we calculated the difference between states of 1/f slope and LZc for each electrode (Figure 3C-F). In a simple linear model, with all animals aggregated and both variables centered (delta LZc and delta 1/f slope), these measures showed a strong inverse relation (adjusted *r^2^* = 0.20, *β* = −2.96; *F*(1, 286) = 71.04; *p* < 0.001), illustrating that electrodes that had a greater modulation by propofol in 1/f slope, also show a greater modulation in their LZc and vice versa. Next, we assessed this effect in one multiple regression model that estimated the linear dependence between these measures for each animal. This model showed a significant main effect (adjusted *r*^2^ = 0.23; *F*(4, 283) = 21.84; *p* < 0.001), and significant individual effects for Monkey 1 (*β* = −3.37; *p* < 0.001), Monkey 2 (*β* = −2.641; *p* < 0.001) and Rat2 (*β* = −6.27, *p* < 0.001), while for Rat1 the linear dependence did not reach significance (*p* = 0.36).

## Discussion

In this article we studied the relation between two apparently dissimilar features of time series from brain field potentials. Our results show a robust and inverse relation between LZc and 1/f slope, constitutive of a one-to-one mapping in both synthetic and empirical data. This relation closely followed an x-inverted asymmetric sigmoid function in the whole range of both measures in iDFT models. This behavior, although scaled, was present even when the spectral power law behavior only comprised a small portion of all frequencies of the signal (Figure 1C, 1D). This is of particular importance as real electrophysiological signals do not show a 1/f spectral power decay in the whole frequency range (44, 52). In a more biophysically constrained model, we observed a similar inverse relation between LZc and 1/f slope, which adjusted to the same mathematical function. We show that this relation follows the balance between excitation and inhibition, with greater complexity and flatter 1/f slopes associated with the predominance of excitatory over inhibitory activity. We probed the link between E/I balance and these two measures in two animal models by directly contrasting 1/f slope and LZc changes due to a pharmacological intervention. Propofol -a GABA agonist-produced changes in both measures consistent with what our models predicted: increased inhibition produced reduced LZc and increased 1/f slope in both monkeys ECoG and rat EEG data. Interestingly, the spatial change by brain state in some electrodes was stronger than others, however, the correspondence of the changes in brain state by propofol showed in both datasets that the electrodes that had a greater change by brain state in 1/f slope, also show a greater change in their complexity. Consistent with prior work (29, 30) we show that changes in E/I balance can be inferred from the 1/f slope of the spectrum. Recently, it has been shown that this can be obtained by modeling excitation and inhibition as disconnected signals (29), as well as in a recurrently connected configuration (30). The disconnected model shows that disruptions in the E/I balance towards inhibition steepens the 1/f slope by increasing the slower time constant associated with inhibitory synaptic currents. The recurrently connected model, however, depends on coupled excitatory and inhibitory dynamics with a realistic biophysical basis that, although more complex, show consistent results with the disconnected model. We show that both models have similar behavior, suggesting that the relation between 1/f slope and LZc -and their predictive power on E/I balance (22)-can be tracked with simple and complex modelling configurations.

Aperiodic neural activity has been studied in a wide range of scientific studies (15) where a divergent set of frequency ranges for the background activity estimates have been used. However, fitting the spectrum across the whole frequency ranges reported in the literature would result in an imprecise fitting. This is consistent with previous work showing the presence of different 1/f slopes at different frequency ranges (30, 42, 45, 52–54), where lower range fittings have a higher correlation with low-frequency oscillations (28) and suggest a note of caution when interpreting 1/f slope results from different frequency ranges as reflecting the same biological mechanism. We have shown, however, that changing the initial and cut-off frequency of the power-law decay does not qualitatively affect the relation between 1/f slope and LZc (Figure 1C, D. Here we analyzed 1/f aperiodic slope as a proxy of background activity and E/I balance, which was most representative in the 30-80 Hz range and is consistent with previous modeling work (29, 30).

While the slope of the spectral power law has been linked to E/I balance (45), LZc reflects the vastness of the repertoire of brain activity patterns (41). Although these two measures may seem unrelated at first, we hypothesize that both reflect a specific type of entropy of brain systems. The entropy of a system can be characterized by the probabilities of each of its possible states (Shannon entropy), but also in terms of the probabilities of the transitions between these states in time, namely its entropy rate (or transition entropy). Low values of 1/f slope represent a flatter power spectrum, which is characteristic of irregular desynchronized brain states, while steeper 1/f slopes showcase mainly low frequency periodic behavior (55, 56). These two extremes can also be characterized in terms of their signals’ transition entropy: flat 1/f slopes (similar to white noise) have low memory and thus high entropy rates, while in mainly periodic signals, its history strongly constrains future values; thus they present low transition entropies. Interestingly, Amigó et al. (57) have shown for electrophysiological signals that LZc closely reflects the entropy rate of the underlying system. This is particularly useful as direct estimations of entropy rate require much longer data series than LZ76 (57, 58). In our implementation of LZc, because we binarize each signal based on its median value, the number of points in each state (ones and zeros) is equal, which results in a constant Shannon or distribution entropy. In this line, we believe signal’s LZc could be reflecting not only the vastness of the repertoire of brain activity, but also specifically the transition entropy of the system. Thus, the strong relation we observe between LZc and 1/f slope suggests both measures are, at least partially, driven by the transition entropy of the underlying brain system.

Future work should include the role of oscillations and phase, as recent evidence has suggested that low frequency 1/f slope is dependent on alpha band activity (53). Despite this potential limitation of our simulations, previous findings have shown that non-linear features -such as LZc-closely relates to the spectral more than the phase component of the signal (58). Consistent with this, we observe the same general behavior in the relation between 1/f and LZc both in EEG and ECoG data, which does present oscillatory activity.

The E/I-balance shapes neurons’ computational properties (59), and therefore behavior and cognition (3). Alterations of this balance have been related to schizophrenia (24), autism (25), and epilepsy (23), which hints it might also play an unexplored role in other neuropsychiatric disorders (26). Moreover, E/I balance is not a static property of the cortex. It changes depending on the behavioral state (60), task demands (60, 61), performance (28) and depending on circadian rhythms (62), which suggests that this property is under fine dynamic control. Recent work have shown brain states and neural complexity can be regulated by ascending arousal activity (63, 64), external stimulation (65) and task demand (66, 67). Future research could address this topic with a multiscale approach to the underlying states of neuromodulation-related psychiatric disorders (68). From this perspective, the readout of E/I balance through brain signal complexity and the power-law of the PSD could be useful for addressing fundamental questions about the modulation of the state dependence of brain computations. This offers new methods to understand the general mechanisms of brain states functioning, as well as broadening the diagnostic and therapeutic tools related to neuropsychiatric disorders.

## Materials and Methods

### Power Spectral Density and 1/f analyses

We employed the same approach to estimate the power-law slope of simulations and monkey ECoG data. We calculated the Power Spectral Density (PSD) by means of Fourier Transforms using Welch’s method as implemented in the MNE toolbox (69). Afterwards, the power-law 1/f slope and offset were obtained using the FOOOF toolbox (14). Aperiodic offset (O) and slope (s) components are obtained by modeling the aperiodic signal according to Equation 1. The FOOOF algorithm decomposes the log power spectra into a summation of narrowband gaussians periodic (oscillations) and aperiodic (offset and slope) components within a broad frequency range. The algorithm iteratively estimates periodic and aperiodic components, removes the periodic ones and estimates again until only the aperiodic components of the signal remain. This allows for estimation of offset and power-law slope with considerable independence from oscillatory behavior, which is particularly important for empirical signal analysis (14, 56). Also, there is evidence revealing spectral “knee”(44) which suggests that one fitting over the whole spectrum will conflate imprecise fitting of the background activity. Previous evidence have shown that an increase in the global network’s inhibition is consistently related to a steepening of the 1/f slope in the range between 30-80 Hz (29, 30). Thus, we use a 30-80 Hz frequency range for all the fittings.

### LZc algorithm

To compute the complexity of time series (both simulated and empirical), we used the LZ76 algorithm as introduced by Lempel and Ziv (34). This algorithm quantifies the number of distinct and non-redundant patterns of a signal and it can serve as a close analogue of the entropy rate of a signal (57, 58). We implemented the LZc algorithm using Python scripts available from previous work (36). Briefly, every time series was first binarized, assigning a value of 1 for each time point with an amplitude greater than the median of the entire signal (5 s), and zero for those below it. Afterwards, the LZ76 algorithm was applied to the resulting so-called symbolic signal. To quantify the number of non-redundant patterns, a sequential evaluation of the signal is performed. At each time point, the algorithm analyzes whether the segment including the following point of the signal can be recreated from the already analyzed signal, be it because it is already present, or because it can be recreated by a simple copying procedure. In this sense, if the following sequence can not be recreated from the previously analyzed signal, then a complexity counter increases. If the next sequence is redundant with respect to the already analyzed signal, the algorithm advances to the next time point without increasing the complexity counter. Once the whole signal is analyzed, the complexity counter (number of non-redundant patterns) is normalized to produce the LZc value, which ranges (asymptotically for long signals) from 0 to 1. A more thorough explanation of the algorithm can be found in the original article (34) and in (36).

### iDFT Model

To study the general relation between powerlaw slope and LZc, we first employed a simple modeling strategy that we denominate iDFT model. We constructed signals with different 1/f slopes, among other spectral parameters, and computed their resulting LZc. Each signal was simulated using 5 seconds of length with 1000 points per second (fs = 1KHz), which resulted in a Nyquist frequency (*Nf*) of 500 Hz. Each time series was initially constructed in the frequency domain as the product of its power and phase components. The power of each frequency component was constructed accordingly to a power-law distribution:

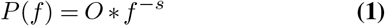

where *P*(*f*) represents power as a function of frequency, *O* is the offset of the curve, the amplitude of the 1 Hz component, and *s* corresponds to the slope of the power-law. Phases were linear in time, starting at an initial phase *θ*_0_ that was randomly assigned from a uniform distribution (–*π* to *π*). iDFT function, as implemented in Numpy (70), was applied to the product of the amplitudes *A*(*f*) (square root of the power) and phase components to obtain the time series data according to Equation 2:

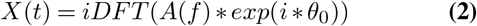

where *X*(*t*) is the resulting time varying signal and i is the imaginary unit. Only positive frequencies were employed. To better model the spectral properties of physiologically plausible neural signals, in addition to constructing signals using the whole range of possible frequencies (0 to *N_f_*) we also applied two types of constraints to the power-law distribution: an initial frequency (*f_0_*; Figure 1C) and a final frequency (*f_f_*; Figure 1D). Both are illustrated in Figure 1A. Specifically, *f*_0_ corresponds to setting all amplitudes of frequencies lower than *f*_0_ to the value of *f*_0_, thus flattening the curve to the left of *f*_0_. On the other hand, applying an *f_f_* corresponds to setting the amplitude of every frequency higher than *f_f_* to zero. To maintain time series stationarity, a requirement of the LZc algorithm (33, 34), all iDFT models were made with a *f*_0_ = 1 Hz unless otherwise stated.

### 1/f slope vs. LZc modeling function

In an attempt to model the observed relation between 1/f slope and LZc, we empirically found that this relation, for a pure power-law iDFT model (Figure 1B), closely followed a particular mathematical behavior:

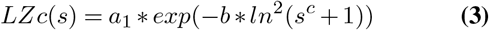

where *s* is the slope of the power-law, *LZc*(*s*) is the LZc as a function of slope and *a*_1_, *b* and *c* are free parameters such that *a_1_* ranges from 0 to 1 and *b* and 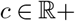. The parameters *b* and *c* modify the shape of the curve, while *a*_1_ is a scaling factor. Without this scaling factor, the image of LZc(s) ranges from (0 to 1), while if *a*_1_ is introduced it ranges from (0 to *a*_1_) without changing the internal structure of the curve. Equation 3 appropriately adjusted pure power-law signals (Figure 1B) and iDFT-data generated with a non-trivial final frequency (*f_f_* < *N_q_*; Figure 1D). Signals with non-trivial *f*_0_ (> 1Hz) did not ranged from 0 to 1 but from a value greater than 0 to 1 (Figure 1C). Because of this, we designed a similar equation that better reflected the required image of the LZc(s) function for non-trivial *f*_0_ cases, introducing a second scaling param-eter *a*_2_:

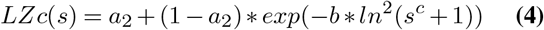

For every fit we employed Equations 3 or 4 using an algorithm that minimized the squares of the differences between data and models as implemented in the *scipy.optimize.curve_fit* function (70). Best fit parameters and *r*^2^ values for goodness of fit for all iDFT simulations can be found in **Table S1** (all *r*^2^ > 0.98). It is important to emphasize that the modeling of the relation between 1/f slope and LZc was conducted by an iterative and empirical approach. Although it is possible that there exists a strict analytical solution for this relation, this escapes the scope of the present article.

### Autocorrelation, time-scale, 1/f and LZc

Given the relation between the general spectral properties of synthetic signals (1/f slope) and LZc, we wondered whether this relation could also be tracked by another non-linear characteristic of time series, namely their time scale (tau). Additionally, there is evidence indicating that tau is related to the initial (or knee) frequency (*f*_0_ in our nomenclature) of the power-law decay of neural data (44). Thus, we calculated the tau of iDFT-generated signals following previously used strategies (71), which are based on the autocorrelation function (ACF). For each time series we first computed its ACF (numpy correlate function), and selected a segment starting at lag = 0 and ending in the first lag point that dropped below 80% of the initial autocorrelation. Within this segment of the ACF, we fitted a quadratic equation and, based on the parameters of this equation, obtained the time lag (tau) at which the ACF reached 50% of its initial magnitude (numpy roots function). Each tau estimation was the average of 20 repetitions.

We analyzed the relation between tau, 1/f slope and *f*_0_ by constructing iDFT models for six 1/f slopes (ranging from 0.3 to 3), each one with 11 values of *f*_0_ (ranging from 1 to 10 Hz). To improve the interpretability of the results, we plotted the multiplicative inverse of *f*_0_, i.e. the period of the slower frequency that follows a power-law behavior as a function of tau (Figure 1E). To analyze the relation between tau and LZc, and its dependence on *f*_0_, we constructed 64 series of iDFT models with different 1/f slopes (ranging from 0.2 to 3.2), each one with five different values of *f*_0_ (0.1, 0.5, 1 and 5Hz). These results are depicted in Figure 1F.

### Conductance-based neural network model and kernel-based LFP estimation

We modeled a standard cortical circuit using a recurrent network of leaky integrate-and-fire (LIF) excitatory and inhibitory neurons with conductance-based synapses. The dynamics of each of the neuron types is described by

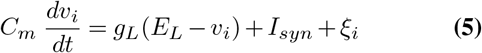

Where *C_m_* represents the membrane capacitance, *v_i_* represents the voltage of neuron *i* and whenever *v_i_* > *v_th_* at time *t*, *v_i_* goes back to the resting membrane voltage *v_rest_* for a refractory period *t_r_*. The synaptic current *I_syn_* received by each neuron i after the spiking activity of all presynaptic neurons *j_i_* described by the following equations

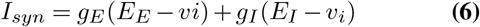

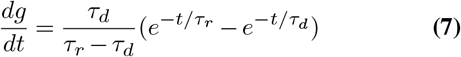

Both excitatory and inhibitory neurons received two types of external inputs (*ξ*) that represent thalamic and cortico-cortical background excitatory drives. Thalamic input was modeled as a Ornstein-Uhlenbeck process and cortico-cortical input as a poisson process. The network structure and parameters were based on the model used in Cavallari et al. (46) according to physiological evidence (Table 1). The network consisted of 5000 neurons, of which 4000 were excitatory and 1000 inhibitory. Neurons forming AMPA-like excitatory synapses and GABA-like inhibitory synapses were randomly connected with a uniform probability of 20%.

**Table 1.** Recurrent LIF model parameters.

| Neuron Type | Parameter Name | Value |
| --- | --- | --- |
| E & I | Resting Membrane Potential | -65 mV |
| E | Population Size | 8000 |
| E | Population Firing Rate | 2 Hz * [0.2, 1, 2, 5, 20] |
| E | Reversal Potential | 0 mV |
| E | Conductance Time Rise | 0.1 ms |
| E | Conductance Time Decay | 2 ms |
| I | Population Size | 2000 |
| I | Population Firing Rate | 5 Hz * [0.5, 2.5, 5, 12.5, 50] |
| I | Reversal Potential | -80 mV |
| I | Conductance Time Rise | 0.5 ms |
| I | Conductance Time Decay | 10 ms |

To compute LFP signals from the recurrent network of spiking neurons, we used a kernel-based method developed by Telenczuk et al. (47). We convolved the spikes of the network with unitary LFP kernels. Changes to E/I balances were simulated as modifications to GABA-like conductances. We introduced a scaling factor that multiplied inhibitory-inhibitory and inhibiotory-excitatory unitary conductances. We employed 21 scaling factors ranging from 1/4 to 4. In this way, we drove the system both towards more excitatory driven situations (low scaling factor) and more inhibition-dominated ones (high scaling factors).

### Macaque ECoG Data

We used an open ECoG database collected from 2 macaque monkeys (Macaca fuscata) during wakefulness, propofol anesthesia (5 and 5.2 mg/kg), and recovery (48). Propofol induced anesthesia was achieved through intravenous propofol injection. Loss of consciousness was defined as the moment when monkeys no longer responded to touch stimuli. The ECoG grid consisted of 128 channels using multichannel ECoG electrode arrays (Unique Medical, Japan). The array was implanted in the subdural space with an interelectrode distance of 5 mm. Electrodes were implanted in the left hemisphere continuously covering frontal, parietal, temporal and occipital lobes. No further preprocessing than the one used by (48) was applied to this data. Since we were interested in assessing differences between brain states during wakefulness and anesthesia and not in the transitions, we only considered periods of closed-eyes wakefulness and anesthesia. We computed LZc and 1/f slope measures of the times series as mentioned above for each electrode, epoch and subject and then averaged LZc and 1/f slope across epochs.

### Rat EEG Data

We used an open EEG database (49) collected from two head - and body - restricted rats during wakefulness and propofol anesthesia (2 mg/kg/min). The EEG recording represents 3 minutes of spontaneous brain activity recorded with 16 EEG channels. All coordinates for electrodes implantation are expressed referring to bregma position, x = medial-lateral axis (–, left hemisphere; +, right hemisphere), y = rostral-caudal axis (–, caudal to bregma; +, rostral to bregma), z = dorsal-ventral axis. Recording electrodes were in contact with the dura and were organized in a grid, symmetric along the sagittal suture, and were placed at the following coordinates (in mm): x = ±1.5, y = +5 (M2); x = ±1.5, y = +2 (M2); x = ±1.5, y = −1 (primary motor cortex; M1); x = ±4.5, y = −1 (primary somatosensory cortex; S1); x = ±1.5, y = −4 (retrosplenial cortex; RS); x = ±4.5, y = −4 (parietal associative cortex; PA); x = ±1.5, y = −7 (secondary visual cortex; V2); x = ±4.5, y = −7 (primary visual cortex; V1); x=0, y = −10 (cerebellum, ground; GND)

### Statistical analyses

Experimental data was visualized using raincloud plots (72). Statistical significance was assessed with a Type-1 error threshold of 0.05. All curve fits were carried out using Scipy optimize function. *R*^2^ was calculated using custom-made scripts. Correlations were assessed by measns of Spearman Correlations. The relation between LZc and 1/f slope in empirical data was evaluated using mixed effects ANOVA. Spatial correspondence between the amount of change in LZc and 1/f slope between resting and propofol anesthesia conditions were assessed by multiple linear regressions.

## ACKNOWLEDGEMENTS

We would like to thank Chile’s national agency for science ANID for their financial support to GB (postdoctoral project N° 3200248), TO (FONDECYT N° 1180932) and to VM (doctoral scholarship N°(21180871). We also thank Fernanda Weinstein for her valuable comments on previous versions of this manuscript.

